# Cerebral Metabolic Changes during Visuomotor Adaptation Assessed using Quantitative FMRI

**DOI:** 10.1101/539130

**Authors:** Catherine Foster, Jessica J Steventon, Daniel Helme, Valentina Tomassini, Richard G. Wise

## Abstract

The neural energetics underlying functional brain plasticity have not been thoroughly investigated in the healthy human brain. A better understanding of the blood flow and metabolism changes underlying plasticity will help us to address pathologies in which plasticity is compromised and, with interventions, could be enhanced for patient benefit.

Calibrated fMRI was conducted in 20 healthy participants during performance of a serial reaction time task which induces rapid motor adaptation. Regions of interest (ROIs) were defined from areas showing linearly decreasing task-induced BOLD and CBF responses. BOLD, CBF and relative CMRO_2_ responses were calculated for each block of the task. The flow-metabolism coupling ratio, *n*, was also calculated for each ROI. Increases from baseline in BOLD, CBF and CMRO_2_ were observed in multiple brain regions including the motor and sensorimotor cortices, cerebellum and hippocampus during SRT task performance, as well as changes in the response amplitude from early to late task blocks reflecting task adaptation. CMRO_2_ responses on average decreased faster than BOLD or CBF responses, potentially due to rapid neural adaptation. However, the mean flow-metabolism coupling ratio was not significantly different between ROIs or across blocks.

Calibrated fMRI can be used to study energetic changes during learning in the healthy brain and could be used to investigate the vascular and metabolic changes underlying reductions in plasticity in ageing and disease.

## Introduction

The brain retains a lifelong ability to adapt through learning and in response to injury or disease related damage, a process known as functional neuroplasticity. Residual neuroplasticity in chronic diseases such as Multiple Sclerosis, or following stroke, can be harnessed in rehabilitation strategies to promote recovery of function. However, the neuronal and vascular mechanisms that promote and limit plasticity are not fully understood. Adequate energy delivery in the form of cerebral blood flow (CBF), which carries oxygen, glucose and other nutrients to tissue, is essential for healthy neuronal function, as is the capacity to metabolise these substrates. In Multiple Sclerosis for example, there is evidence of both CBF (D’haeseleer, Cambron, Vanopdenbosch, & De Keyser, 2011; Ota et al., 2013) and metabolic dysfunction (Fan, Evans, Stout, Rosen, & Adalsteinsson, 2015; Ge et al., 2012; Kidd et al., 1999) which may play a central role in limiting plasticity. The neural energetics underlying functional brain plasticity have not been thoroughly investigated experimentally in the healthy human brain. A better understanding of the blood flow and metabolism changes which occur during motor skill acquisition, and which facilitate plasticity, is needed before characterisation in disease, and subsequent translation to informing treatment interventions to maintain or recover function.

Calibrated fMRI enables measurement of regional CBF and relative changes in the rate of cerebral metabolic oxygen consumption (CMRO_2_) through the addition of hypercapnic calibration (Davis, Kwong, Weisskoff, & Rosen, 1998; Hoge et al., 1999) during dual-acquisition of BOLD and CBF weighted images. The technique has potential applications in identifying clinically relevant abnormalities in vascular and metabolic function which are not visible using BOLD fMRI alone. CBF and CMRO_2_ provide additional information which aids interpretation of fMRI studies of ageing and disease where neurovascular coupling (NVC) is likely to be altered (Restom, Bangen, Bondi, Perthen, & Liu, 2008). For example, greater BOLD responses with increasing age during a Stroop task have been reported, alongside a reduced CMRO_2_ increase in response to the task (Mohtasib et al., 2012). This suggests that as CBF was unaffected by age in this cohort, the greater BOLD signal changes were due to a reduction in the CMRO_2_ response. As CMRO_2_ and neuronal firing are closely coupled, a decreased neural response with age is the most likely explanation for these results. Such changes in vascular reserve and NVC would not have been evident with BOLD alone, and the results demonstrate the value of calibrated fMRI in any study where cerebral energetics may be altered by experimental conditions or over time.

CMRO_2_ changes during the task may also reflect increases in aerobic glycolysis which have been suggested during learning, and motor adaptation (Goyal, Hawrylycz, Miller, Snyder, & Raichle, 2015). Recently, using PET, Brodmann’s area 44 (BA44) showed increases in aerobic glycolysis and reductions in O_2_ metabolism following a visuomotor adaptation task (Shannon, Neil, Vlassenko, Shimony, & Rutlin, 2016), the latter being associated with reductions in synaptic activity. The sustained aerobic glycolysis increases in BA44 were interpreted as a local metabolic change resulting from functional remodelling following learning. As aerobic glycolysis varies regionally (Vaishnavi, Vlassenko, Rundle, Snyder, & Mintun, 2010), it is therefore likely that CBF and CMRO_2_ responses will also differ within task networks. In addition, Madsen et al. (1995) measured global CBF, glucose, and O_2_ consumption during the Wisconsin Card Sorting Task using the Kety-Schmidt technique. During the task, CBF and glucose consumption increased but O_2_ consumption did not. The observed increase in glucose consumption showed that there was an increase in aerobic glycolysis, which persisted for 40 minutes after task completion, although CBF levels returned to baseline. However, the Kety-Schmidt technique is global, and any localised O_2_ consumption changes would not have been detected. Nevertheless, these results suggest that aerobic glycolysis plays an important role in long term depression (LTD) and long term potentiation (LTP) processes which occur with learning and plasticity. The role of aerobic glycolysis in learning and skill acquisition must be taken into consideration when interpreting CMRO_2_ changes as aerobic glycolysis changes may affect the net change in CMRO_2_ during such tasks (Vaishnavi et al., 2010).

The aim of the current study was to measure BOLD, CBF and CMRO_2_ responses during performance of a serial reaction time (SRT) task (Nissen & Bullemer, 1987) using calibrated fMRI. In the SRT task, participants respond to a sequence of stimuli that appear one-by-one at various locations on a screen. Participants respond by indicating the current stimulus location which follows a repeating pattern (see figure 1) allowing participants to identify this sequence with practice, improving the accuracy and speed of responses. Visuo-motor task performance can be improved over short periods of time, accompanied by haemodynamic changes in task-relevant regions, which are thought to reflect short-term plasticity in the adult brain (Fernández-Seara, Aznárez-Sanado, Mengual, Loayza, & Pastor, 2009; Olson et al., 2006; Shannon et al., 2016). Previous work has demonstrated BOLD and CBF task responses in motor and visual cortex as well as prefrontal regions and the cerebellum during task performance (Ungerleider, Doyon, & Karni, 2002). BOLD signal reductions over time, due to task adaptation have also been observed within a single MRI session (Floyer-Lea & Matthews, 2004; Shannon et al., 2016). Therefore, we expected to observe BOLD signal decreases in task-relevant areas as motor adaptation occurred. CBF and CMRO_2_ may change dynamically with adaptation to the task in ways that are not visible by looking solely at the BOLD response as it is the result of changes in vascular and metabolic processes. For example, although BOLD and CBF responses have been reported during motor skill learning task (Olson et al., 2006), it is not established whether their changes over time follow similar patterns or whether there are alterations in the CMRO_2_ response, as previously no changes (Madsen et al., 1995) and reductions in CMRO_2_ (Shannon et al., 2016) have been reported during skill learning.

**Figure 1.**
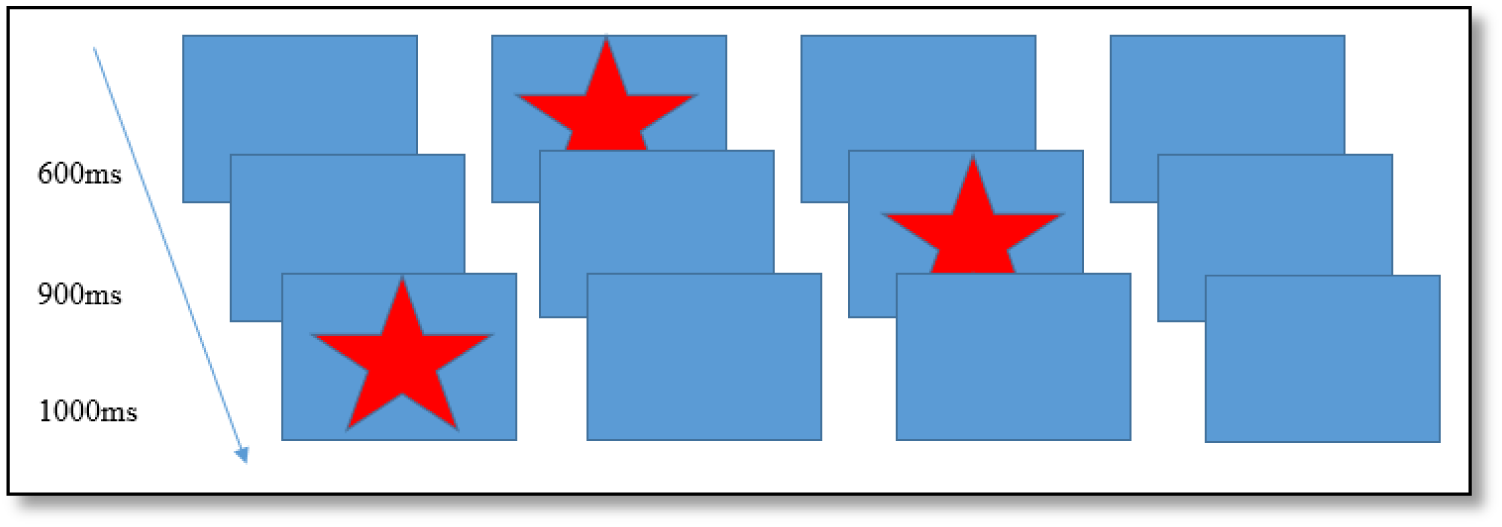
Schematic of the SRT presentation, ISIs on the left indicate example times between trials.

To examine motor task adaptation, BOLD, CBF and CMRO_2_ responses were calculated for each block of an SRT task in regions which showed reducing BOLD and CBF responses across the task. Significant changes in each parameter from baseline were investigated along with flow-metabolism coupling differences between brain regions and task blocks. Lastly, regression analysis was conducted to determine whether CBF, CMRO_2_ or BOLD predicted task performance.

## Methods

### Participants

20 right handed, healthy participants (10 female, mean age 25±4.6) took part in this study. All participants were non-smokers and educated to university level. The study was approved by the Cardiff University School of Psychology Research Ethics Committee and performed in accordance with the guidelines stated in the Cardiff University Research Framework (version 4.0, 2010). Informed written consent was obtained for all subjects.

### Imaging

Imaging was performed on a whole body 3T MRI (GE Excite HDx, Milwaukee WI, USA) system using an 8-channel receive-only head coil. Simultaneous perfusion and BOLD weighted data were acquired with a single subtraction PICORE QUIPSS II (Wong, Buxton, & Frank, 1998) pulsed arterial spin labelling (PASL) sequence (non-commercial) with a dual-echo gradient-echo readout(Liu, Wong, Frank, & Buxton, 2002) and spiral k-space acquisition (Glover, 1999).

Imaging parameters for functional scans (task and hypercapnic calibration) were: TR = 2.4s, TE1 = 2.7ms, TE2 = 29ms, TI1 = 700ms, TI2 = 1.5s (most proximal slice), FOV = 19.8cm, flip angle = 90° matrix size = 64×64, slice thickness 7mm, 1.5 mm gap, 3.1mm in plane resolution with 15 slices. Label thickness was 200mm with a 10mm gap between the end of the label and the most proximal imaging slice. A separate single volume M_0_ scan was acquired using the same parameters, except TR=4s, to measure the equilibrium brain tissue magnetisation of cerebrospinal fluid (CSF) for absolute CBF estimation. For registration, a 3D T1-weighted fast spoiled gradient echo sequence was acquired; TR=7.9ms, TE=3ms, 256 x 256, slice thickness = 1 mm, giving a resolution of 1mm^3^.

Physiological monitoring was performed using a respiratory belt placed just below the ribs to monitor ventilation and a pulse oximeter to obtain cardiac traces. A sampling line connected to a tightly fitted face mask (Quadralite Intersurgical, Wokingham, Berkshire, UK) was used to record expired P_ET_CO_2_ and P_ET_O_2_ concentrations using the Biopac system (Biopac®, Worcestershire, UK). The face mask was connected to a breathing circuit used to deliver gas mixtures and followed the design of (Tancredi, Lajoie, & Hoge, 2014). The MEDRAD system (MEDRAD, Pittsburgh, PA) was used to monitor blood O_2_ saturation during hypercapnia.

### Visuomotor Task

The SRT (Antonio, Case, Harman, & Lacey, 1987) is a visuomotor task which has been used previously in fMRI studies in healthy subjects as well as in patient groups such as MS, chronic stroke and Huntington’s Disease (HD) (Bonzano et al., 2011; Boyd & Winstein, 2001; Knopman & Nissen, 1991).

A modified version of the SRT developed by (Nissen & Bullemer, 1987) was used as the visuo-motor learning task during imaging acquisition. The task was projected via a screen inside the scanner at a frame rate of 60Hz and a resolution of 1024×768. A star appeared on the screen in a sequence of 4 boxes (figure 1), participants responded by pressing the corresponding button on a button box in their right hand. The 12-minute task consisted of 6 blocks of a 12-item sequence repeated 6 times with variable inter-stimulus interval (600-1000ms) interspersed with 3 blocks of a pseudorandom sequence to assess response latency decreases related to task familiarisation rather than sequence learning. Participants were not informed that there was a repeating sequence during task instructions.

### Hypercapnic Calibration

The SRT task was followed by hypercapnic calibration to obtain a measure of cerebrovascular reactivity (CVR) to CO_2_ for estimation of CMRO_2_. Participants breathed through a tight-fitting face-mask as described above, gases were administered from gas cylinders connected to an in-house built manually controlled flow meter system. Gases were piped through a mixing chamber with three feeding lines coming in for the delivery of medical air, 5% CO_2_, and medical oxygen. Medical oxygen was not administered but was connected in case of emergency. The scan began with a 2-minute normocapnia period during which participants breathed medical air (20.9% O_2_ balance N2) with a flow rate of 30 L/min. This was followed by a rapid switch to 2 minutes of hypercapnia where an increase in P_ET_CO_2_ of +7 mmHg was targeted. In total, the scan consisted of three two-minute blocks of normocapnia and two two-minute blocks of hypercapnia.

## Data Analysis

### Image Preprocessing

Perfusion and BOLD weighted images were created from the first and second echo data respectively. Physiological noise correction was carried out using a modified RETROICOR technique (Glover, Li, & Ress, 2000) to remove cardiac and respiratory noise components from the BOLD and CBF task data. First and second harmonics of the respiratory and cardiac cycle along with the interaction term were regressed from the raw CBF signal (before tag and control subtraction) in a GLM framework. In addition, variability related to CO_2_, O_2_, respiration and heart rate were removed from the SRT task run (Birn, Murphy, Handwerker, & Bandettini, 2009). Surround averaging was applied to the BOLD weighted images to remove contamination from perfusion weighting (Liu & Wong, 2005). Perfusion signal modelling was carried out within FEAT (FMRI Expert Analysis Tool) to model the difference between control and tag images in the timeseries.

### Task Response Modelling

One subject was excluded from the analysis due to a low task response rate. The remaining 19 subjects’ BOLD and CBF task responses were analysed using a general linear model (GLM) within FEAT (FMRIB’s Software Library, www.fmrib.ox.ac.uk/fsl) with high pass filtering (cut off 80s). The voxelwise GLM was used to identify statistically significant BOLD and CBF responses across all sequence blocks. FEAT contrasts were set up to investigate positive and negative task vs. rest activity. Further contrasts modelled linear response changes over time, using the task timing regressor to model the average change across the experiment as opposed to a block by block change. Cluster based thresholding was applied to define significant BOLD and CBF task responses (z > 2.3, p < 0.05, FWE corrected). A second FEAT GLM was carried out with contrasts set to calculate BOLD and CBF responses for each individual sequence and random block.

Individual subject’s functional data were registered to the high resolution T1-weighted structural image using FLIRT, FMRIB’s linear image registration tool (Jenkinson & Smith, 2001), with 6 degrees of freedom. The high-resolution images were then registered to the Montreal Neurological Institute (MNI) standard space with 12 degrees of freedom. FEAT contrasts were set up to investigate positive and negative task vs. rest activity. All subjects’ data were then entered into a higher-level FEAT analysis to define functional regions of interest (ROIs) for further analysis.

### Definition of ROIs

ROIs were created from regions where there was a linearly decreasing component to the task-induced-signal (BOLD and CBF) over sequence blocks, to investigate training related adaptation. All reported ROIs were created from the intersection between BOLD and CBF task responses unless otherwise stated. Random blocks were not included in ROI creation. These areas were then separated into anatomical regions using the Harvard-Oxford cortical and subcortical atlases within FSL. The thalamus and BA44 were also included as ROIs due to the previously reported involvement of these regions in motor learning (Hardwick, Rottschy, Miall, & Eickhoff, 2013; Shannon et al., 2016). However, BA44 did not show a significant mean BOLD or CBF signal decrease across the task and in the thalamus, only a BOLD signal decrease was observed. For each ROI, the parameter estimate (PE) for each stimulus block was used to calculate the percentage CBF change or BOLD response for each stimulus block.

### Calculation of CVR and CMRO_2_

CVR to CO_2_ was calculated according to the method described previously (Bright & Murphy, 2013), where the beta weight calculated for the CO_2_ regressor reflects the percentage BOLD or CBF signal change caused by hypercapnia and is normalised by the change in in end-tidal CO_2_ (mmHg). CVR was calculated using the voxel-averaged timeseries in each ROI. As the timing of the haemodynamic response is not uniform across the brain and there are delays in the physiological response to CO_2_, the end-tidal CO_2_ regressors were selected based on which of the 97 time shifts applied produced the best fit to the data.

The Davis model (Davis, Kwong, Weisskoff, & Rosen, 1998) was used to calculate CMRO_2_ from normalised BOLD and CBF data in each ROI. The hypercapnia measurement was performed to estimate the scaling parameter *M* (see equation 2A) which represents the estimated maximum BOLD signal response upon washout of all deoxyhaemoglobin according to the calibrated fMRI equation (Davis et al., 1998) (equation 2B). This model assumes that the targeted level of hypercapnia does not change CMRO_2_. The values for α and β must also be assumed. In the model, α represents the change in CBV as a function of CBF, the original value of this exponent as proposed by Grubb (1974) was 0.38 to describe arterio-venous blood volume. More recently it has been established that the volume of the deoxyhaemoglobin compartment, venous CBV, is what is required to calculate *M*. Chen, & Pike (2010) used steady-state flow and volume changes to estimate the power–law relationship between CBV and CBF and the model fit produced a coefficient of 0.18 which is comparable to simulation work (Griffeth & Buxton, 2011) and values between 0.18-0.23 and are more commonly used at 3T. However, being a biological parameter, actual α values are likely to vary with age, health status and under different experimental conditions.

The parameter β which equals 1.5 in the original equation is a constant representing the relationship between blood oxygenation and the BOLD signal. As relaxivity is field dependent an optimised value of 1.3 tends to be used at 3T (Bulte et al., 2012; Mark, Fisher, & Pike, 2011). Values of α = 0.20 β =1.3 were used in this study (Bulte et al., 2012).

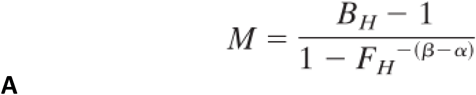

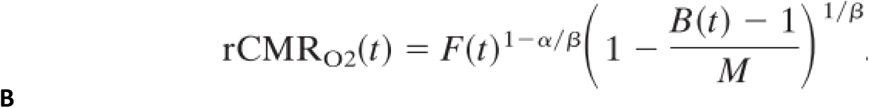

Equation 1 (A) Equations for the calculation of *M* (A) and relative CMRO_2_ (rCMRO_2_) (Davis et al., 1998) (B). B_H_ and F_H_ are BOLD and CBF CVR measures, α is the estimated ratio of fractional change in CBF to CBV and β which represents the field strength dependent relationship between blood oxygenation and the BOLD signal, F(t) = CBF at time (t), B(t) = BOLD at time (t).

### Statistical Analysis

Statistical analyses were carried out using R (https://www.R-project.org) and SPSS version 20.0 (IBM Corp., Armonk, N.Y., USA). The false discovery rate (FDR) was used correct for multiple corrections where such corrections were necessary; a corrected p-value of < 0.05 was considered significant in all cases. To derive an overview of block-by-block responses, repeated measures ANOVAs with tests for sphericity were used to compare each block to a zero baseline, and to compare responses in sequence blocks 1, 2 and 6 against each other.

Repeated measures ANOVAs were used to investigate changes in the flow-metabolism coupling ratio *n*, the ratio of the fractional change in CBF relative to the fractional change in CMRO_2_ (Buxton, Uludağ, Dubowitz, & Liu, 2004), in each ROI and between blocks 1, 2 and 6. Repeated measures ANOVAs were again conducted for behavioural data to identify significant RT changes from sequence block 1 vs. all subsequent sequence and random blocks. RT was then used to investigate relationships between behaviour and neurophysiological data. Finally, multiple linear regression analysis was used to investigate the relationships between RT and imaging-based responses.

## Results

### Behavioural Responses

A one-way repeated measures ANOVA was carried out to compare performance across sequence blocks. Mauchley’s test for sphericity was significant with p < 0.001, therefore degrees of freedom were corrected using Greenhouse-Geisser estimates of sphericity, ε = .441. The results showed that there was a significant effect of time on performance; response accuracy improved over time; *F*(2.2, 38.3) = 12.47, p < 0.001. Follow up comparisons indicated that all subsequent blocks had a significantly higher accuracy rate than block one. There was a 16.9±9% accuracy improvement from sequence block 1 to sequence block 6, p < 0.001. Accuracy was higher in sequence blocks than random blocks, apart from sequence block 1 which had the lowest performance of all blocks (Figure 2a). Random blocks 1 and 2 had significantly lower accuracy than sequence blocks 3-6. However, random block 3 scores were only significantly lower than sequence blocks 3, 4 and 6, all p < 0.05. This shows that performance improvements over time can largely be attributed to sequence learning rather than more general motor skill improvements.

**Figure 2.**
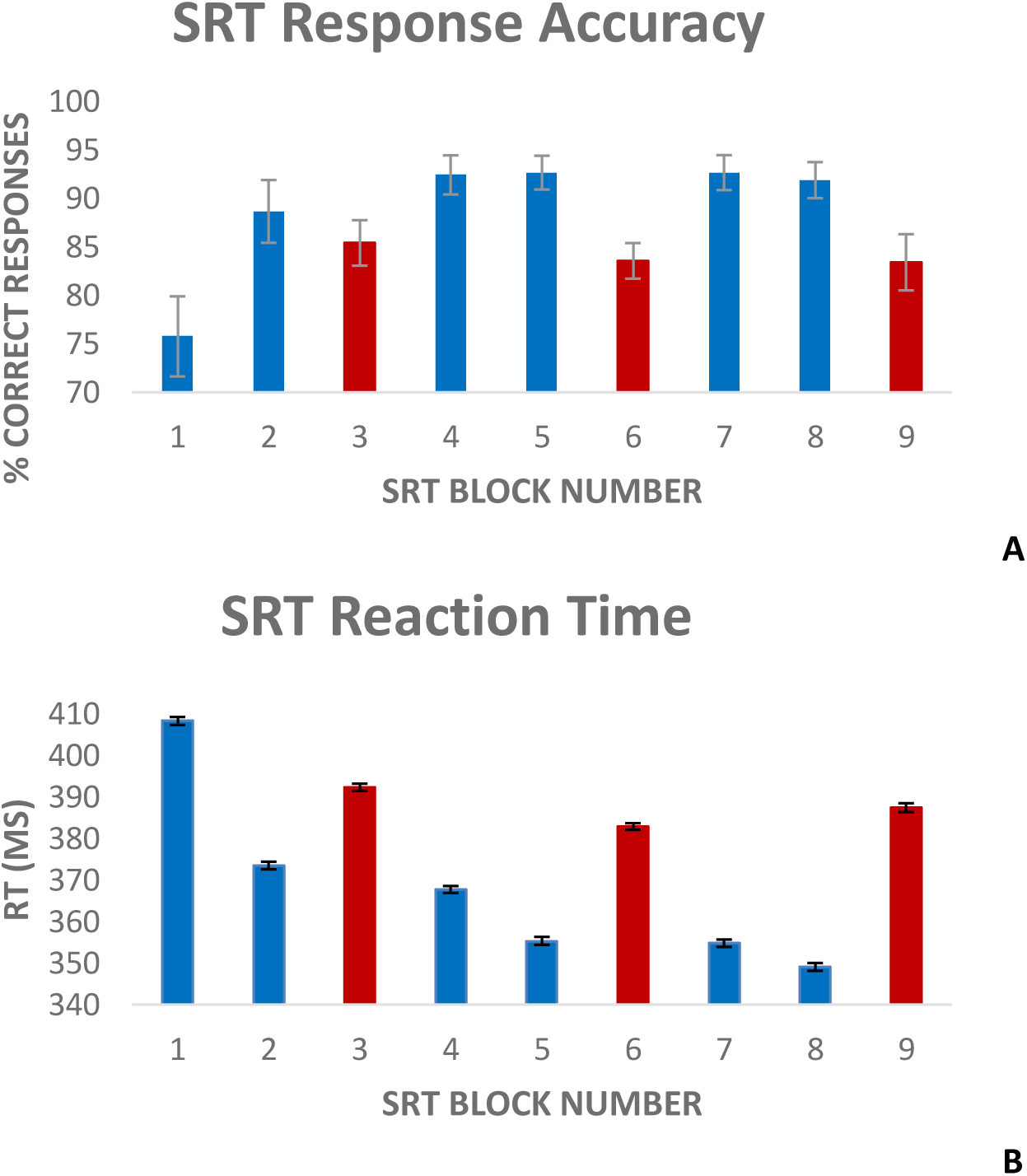
Average (mean ± SEM) response accuracy per block (A), and average response latency (B) per task block. Error bars represent the standard error of the mean across participants. Blocks 3, 6 and 9 (shown in red) represent pseudorandom sequence blocks.

There was a significant effect of time on reaction speed; *F*(5, 85) = 298, p < 0.001. Follow up comparisons showed that RT decreased by 5.1±11% from block 1 to block 6, p < 0.001. Random blocks had a greater response latency, again except for sequence block 1 (Figure 2B). Reaction time (RT) was significantly longer in random block 1 than sequence blocks 2-6, random block 2, RT in random blocks 2 and 3 was longer than sequence blocks 3-6, all p < 0.05.

### Imaging Data

#### Overview

Figure 3a-f shows the mean BOLD and CBF task responses, and regions where signal decreased on average across task blocks, respectively as well as the conjunction between BOLD and CBF responses in each case. There were no areas of statistically significant CBF, or BOLD response increases from rest across the task. Table 1 shows the group mean CVR and *M* values for each ROI shown in figures 4 and 5.

**Table 1.** Mean (SEM) CVR and *M* values for each ROI shown in figures 4 and 5A-K.

| **ROI** | **BOLD**<br>**CVR**<br>**%/mmHg** | **CBF CVR**<br>**%/mmHg** | ***M* %** |
| --- | --- | --- | --- |
| **Global Signal**<br>**reduction**<br>**across the**<br>**task** | 0.17<br>(0.09) | 2.4 (1) | 8<br>(0.5) |
| **ACC** | 0.24<br>(0.02) | 2.1 (0.2) | 13<br>(2.5) |
| **BA44** | 0.23<br>(0.03) | 2 (0.2) | 15<br>(3.3) |
| **Brainstem** | 0.22<br>(0.02) | 2.6 (0.3) | 8 (1) |
| **Cerebellum** | 0.23<br>(0.04) | 2.5 (0.3) | 8<br>(1.7) |
| **Hippocampus** | 0.11<br>(0.02) | 1.9 (0.2) | 9.5<br>(2.8) |
| **Insula** | 0.16<br>(0.01) | 2.1 (0.2) | 4 (3) |
| **LOC** | 0.12<br>(0.01) | 2.2 (0.3) | 4.5<br>(1.7) |
| **M1** | 0.2 (0.02) | 2.3 (0.3) | 13<br>(3.6) |
| **S1** | 0.2 (0.03) | 2.2 (0.3) | 17<br>(5.6) |
| **Precuneus** | 0.3 (0.04) | 2.1 (0.3) | 13<br>(2.7) |
| **Thalamus** | 0.45<br>(0.05) | 3 (0.4) | 19<br>(4.2) |

**Figure 3.**
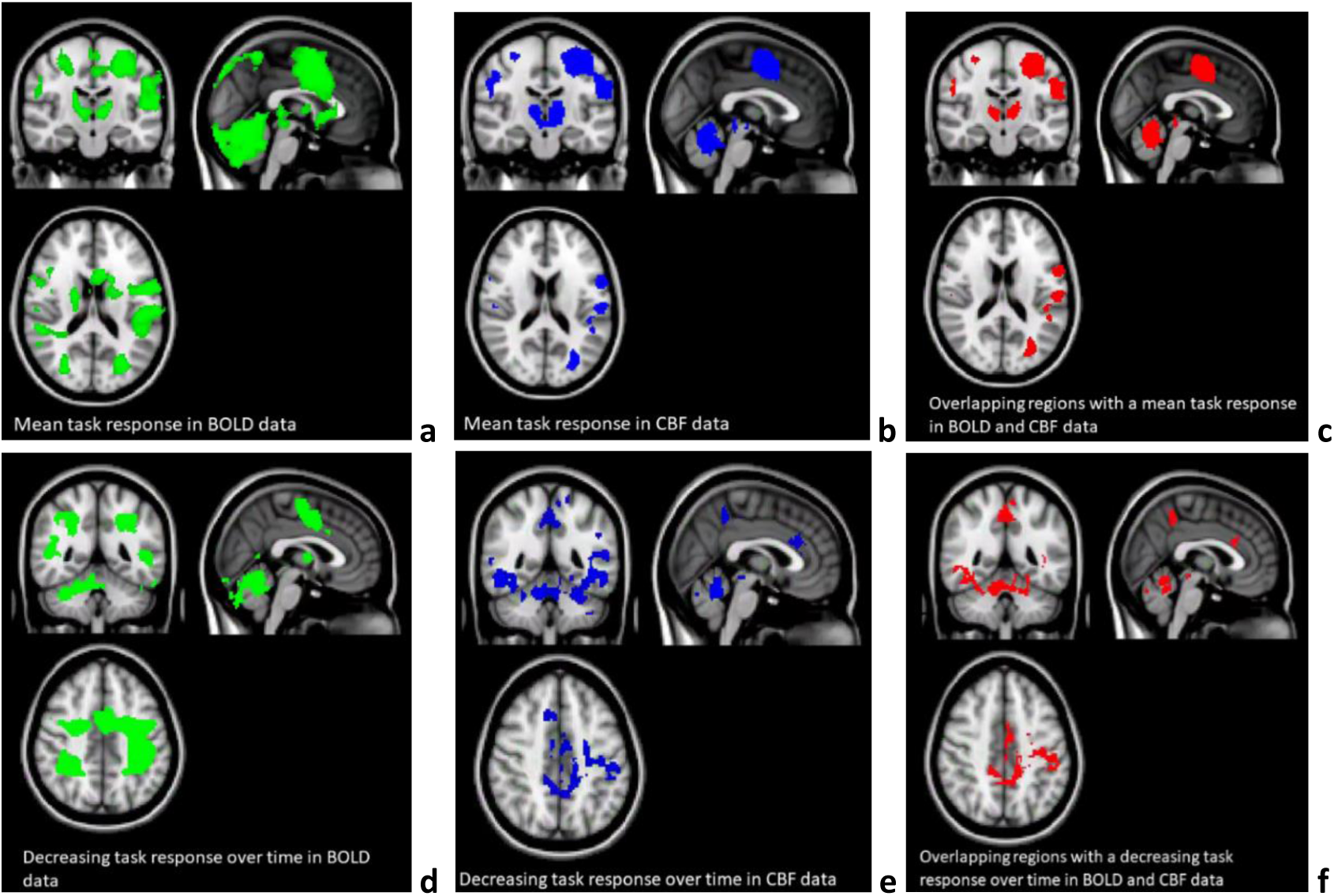
Maps (a-c) show the mean BOLD signal response to the task (green), the CBF response (blue) and regions with overlapping BOLD and CBF task-related signal increases from rest (red) (see design column 3 in supplementary figure 1). Random blocks were not included in this analysis. Maps (d-f) show areas of BOLD signal decrease over time during the SRT task (green), decreasing CBF responses (blue) and overlapping BOLD and CBF signal decreases across the task (see design column 7 in supplementary figure 1). As above, random blocks were not included in this analysis.

**Figure 4.**
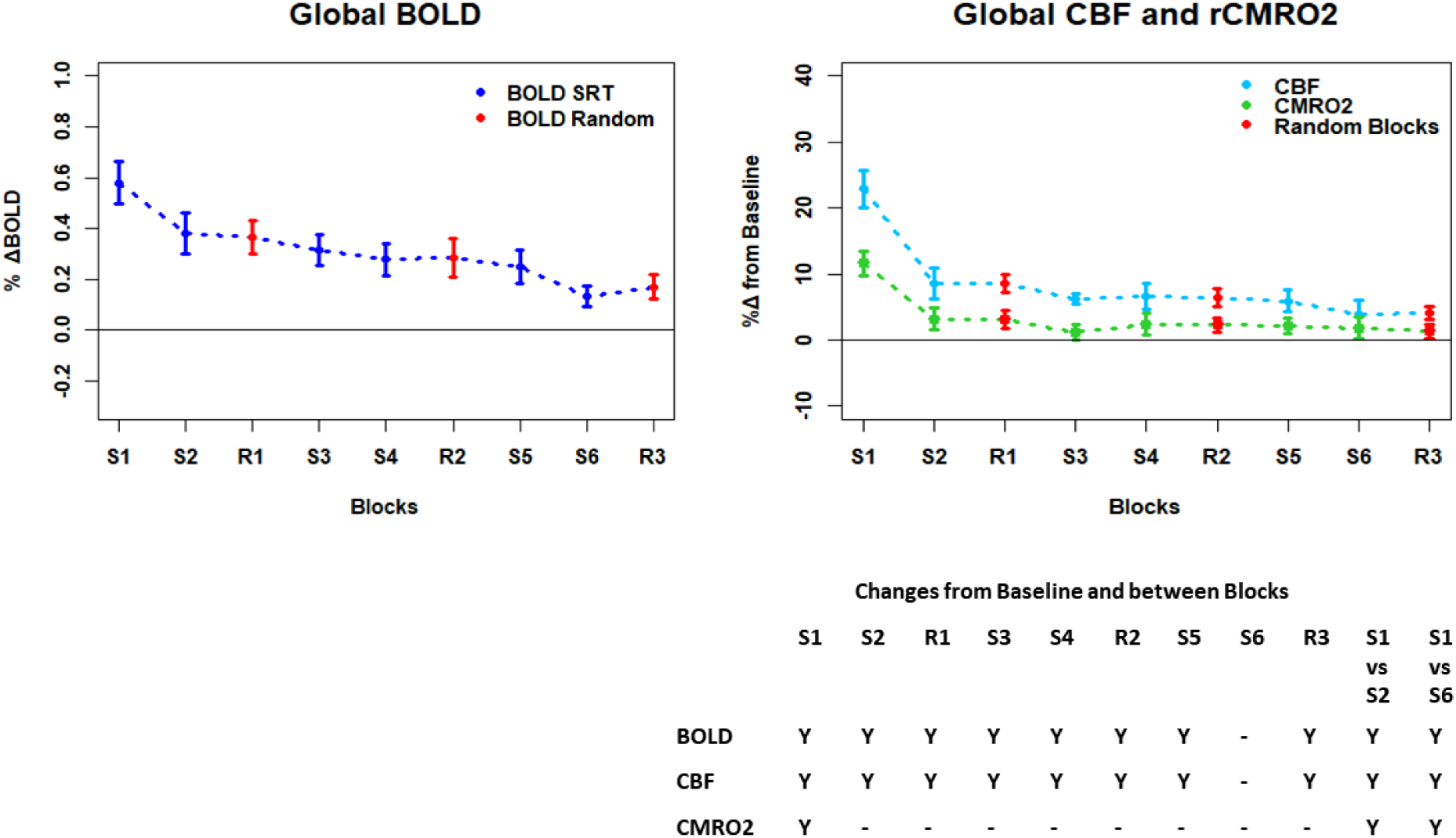
Plots showing the mean ± SEM responses for each task block in the global mean reduction ROI shown in figure 3f. S1 = SRT block 1, R1 = random block 1. Y = Yes to indicate statistically significant (p < 0.05), signal changes from baseline and between first and second and first and last SRT blocks.

**Figure 5.**
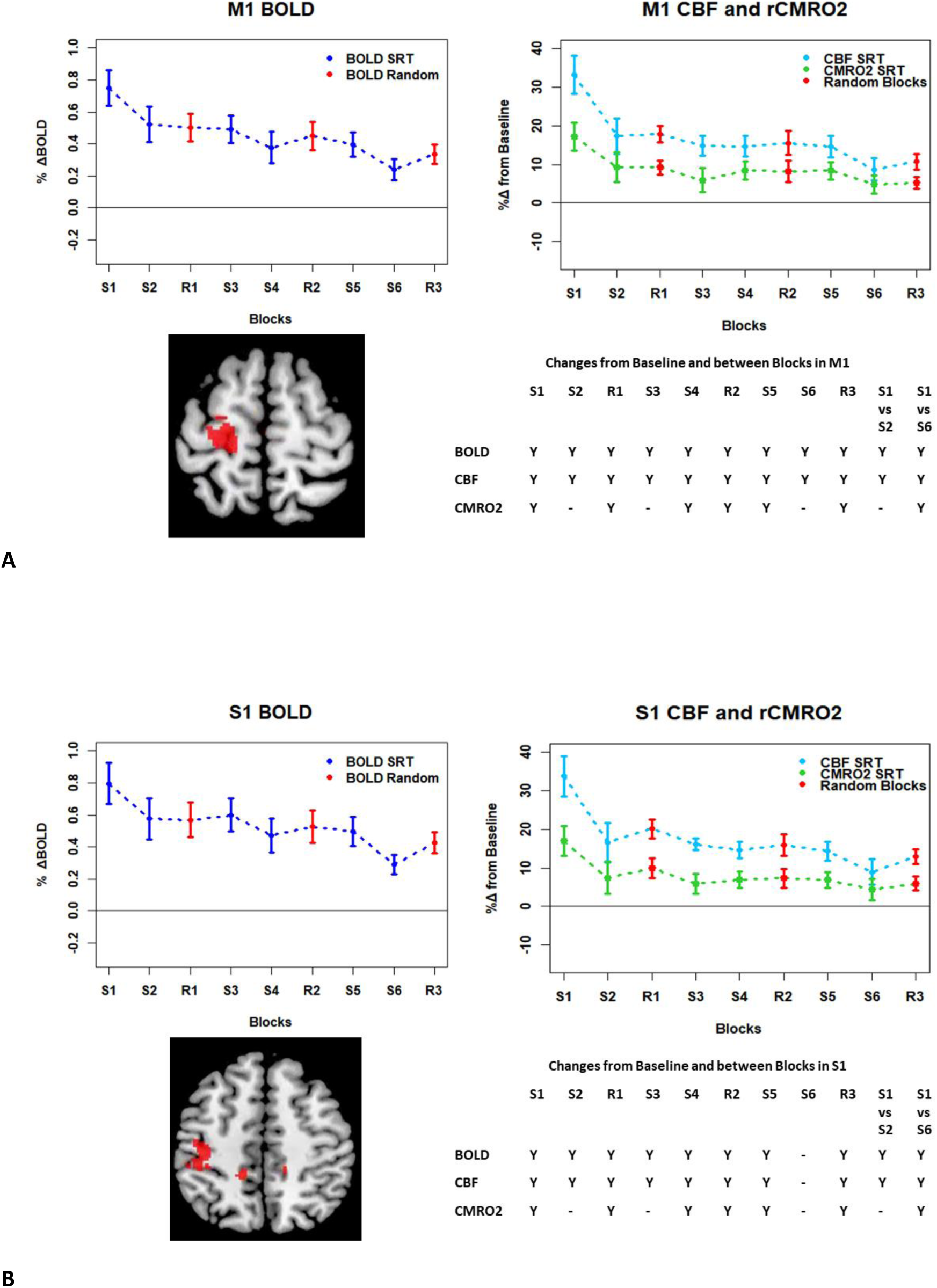
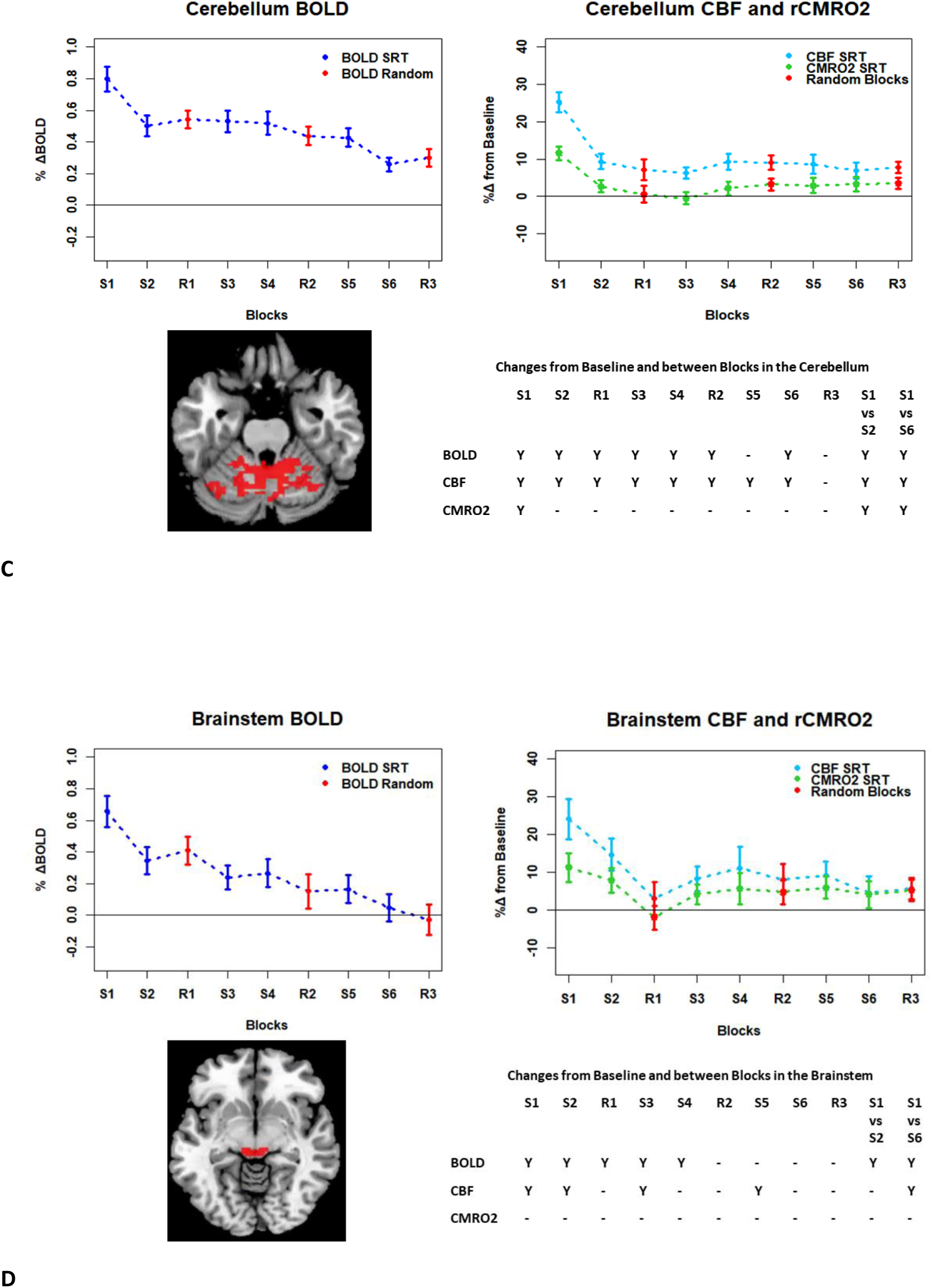
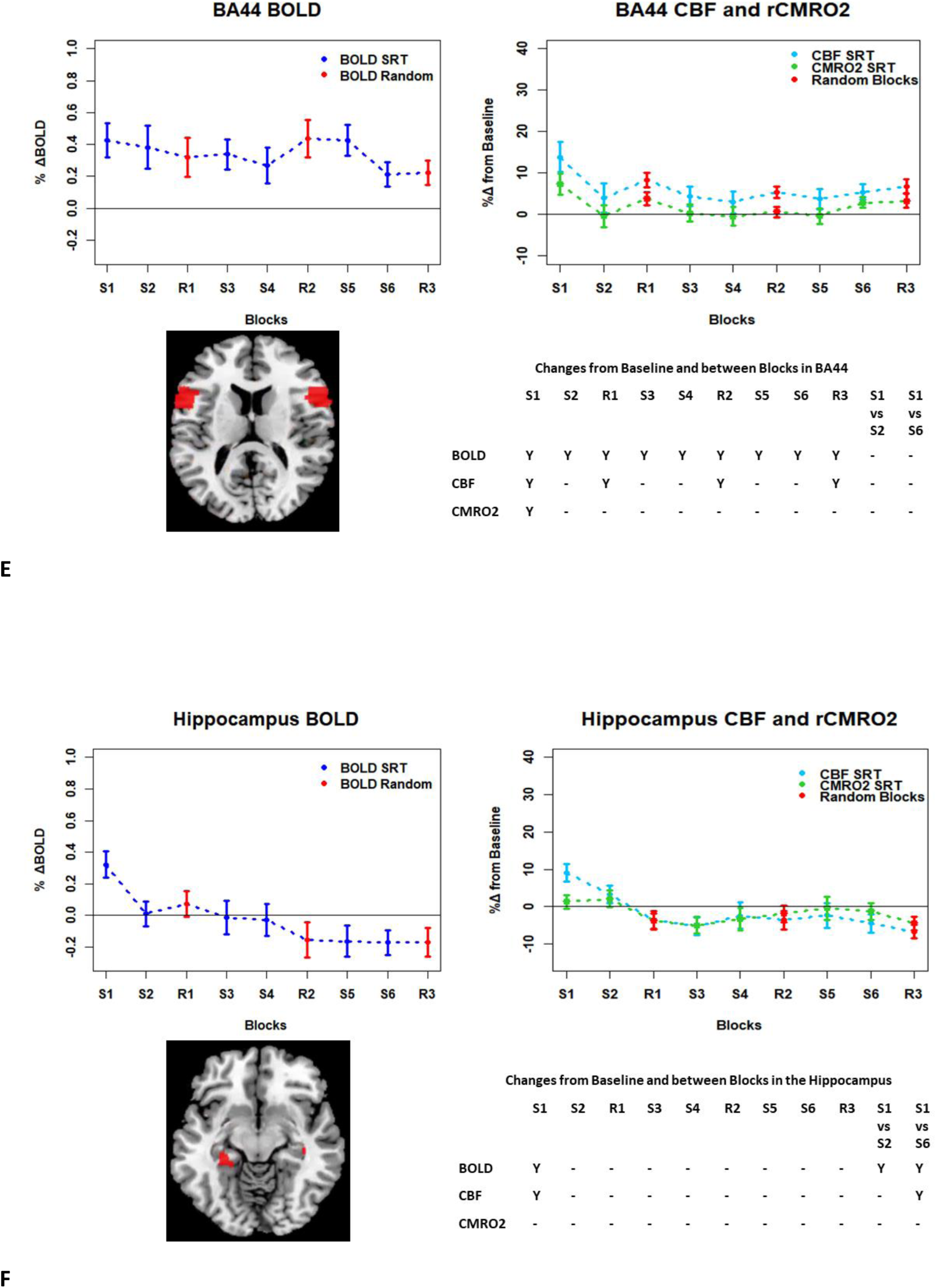
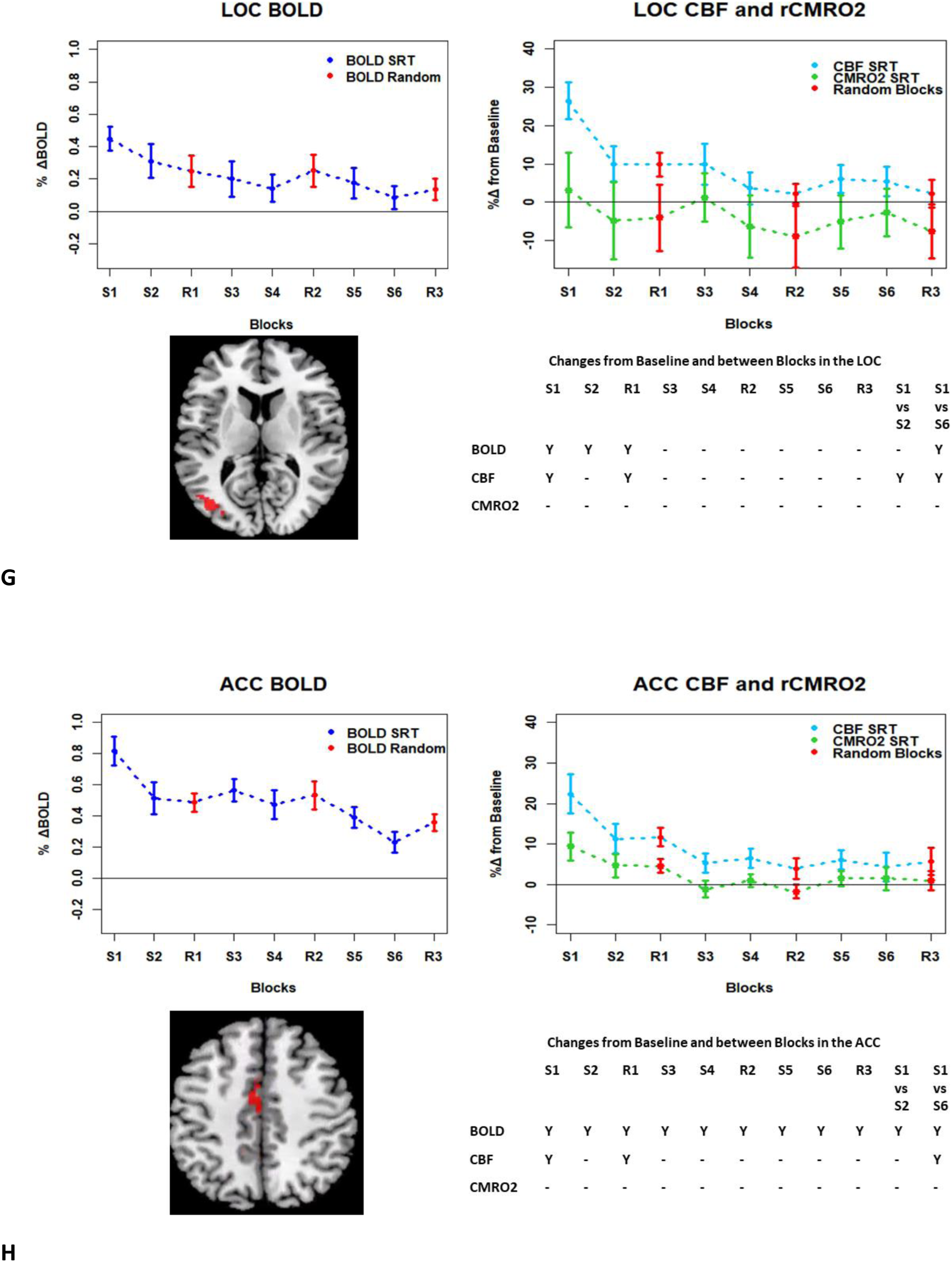
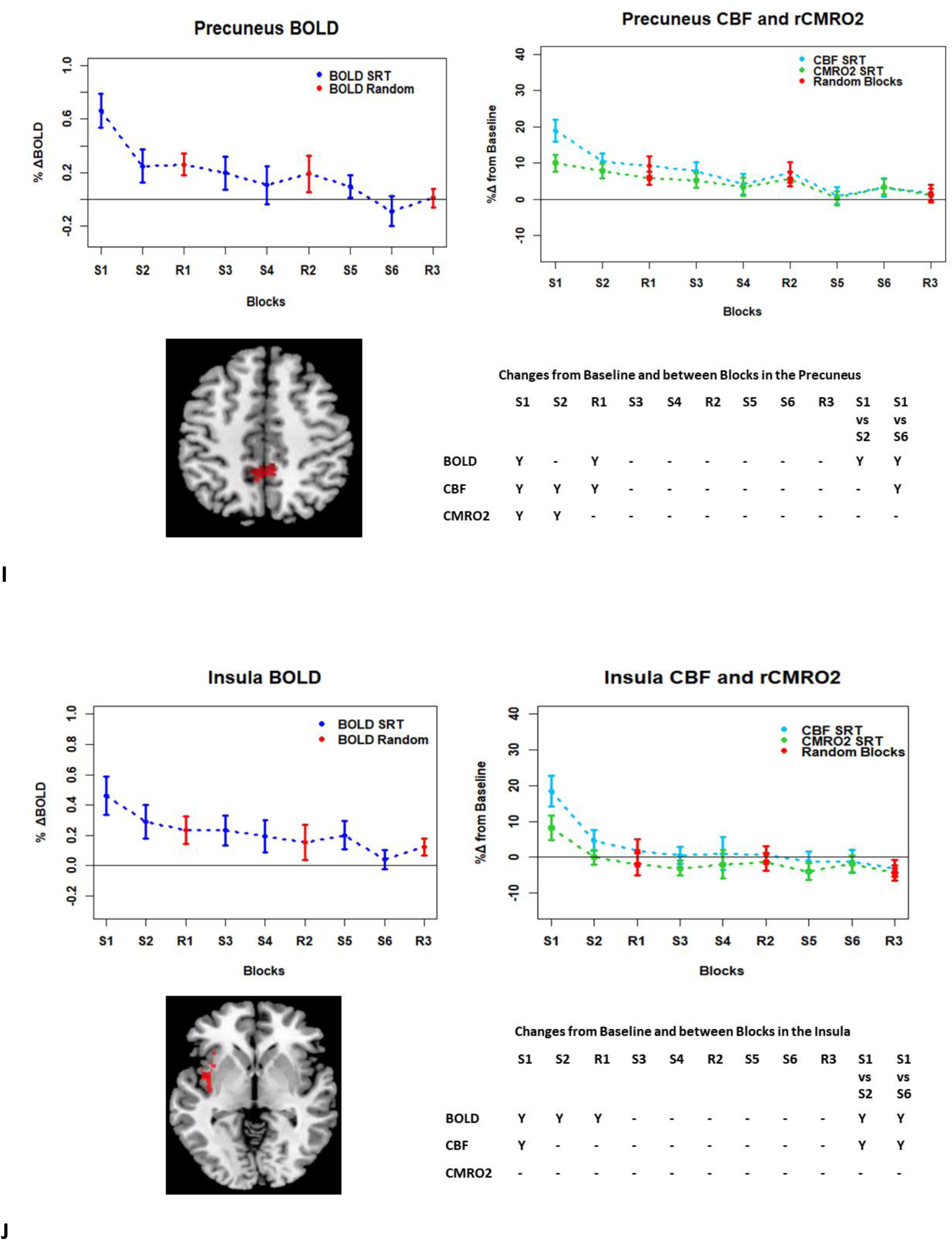
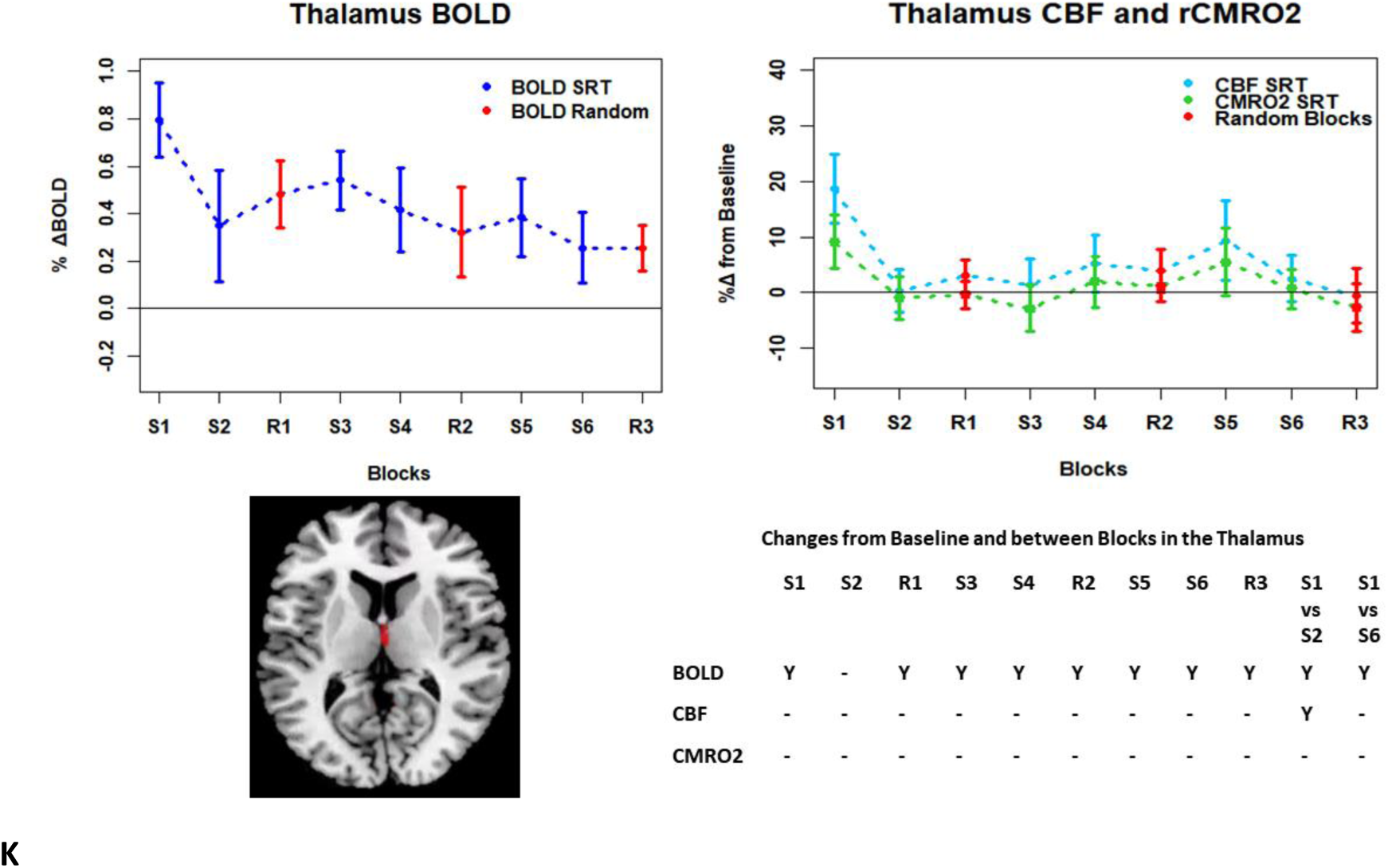
Plots showing the mean ± SEM responses for each task block in different anatomical regions showing a mean (BOLD and CBF) signal reduction across the task. S1 = SRT block 1, R1 = random block 1. Y = Yes to indicate statistically significant (p < 0.05), signal changes from baseline and between first and second and first and last SRT blocks.

### Anatomical ROIs: BOLD, CBF and CMRO_2_ Responses in Each Task Block

ROIs defined from the intersection between BOLD and CBF task response reductions (shown in figure 3f) and subdivided into 9 anatomical areas were investigated: the precentral gyrus (M1), the postcentral gyrus (S1), insula, lateral occipital cortex (LOC), precuneus, anterior cingulate cortex (ACC) hippocampus, brainstem and cerebellum. For the additional ROIs, the thalamus showed a significant BOLD signal reduction over time, and BA44 did not show any signal reduction. Figures 5 and 6 show the mean BOLD, CBF and CMRO_2_ responses for each task block across the global reduction ROI and for each of the anatomical ROIs respectively.

**Figure 6.**
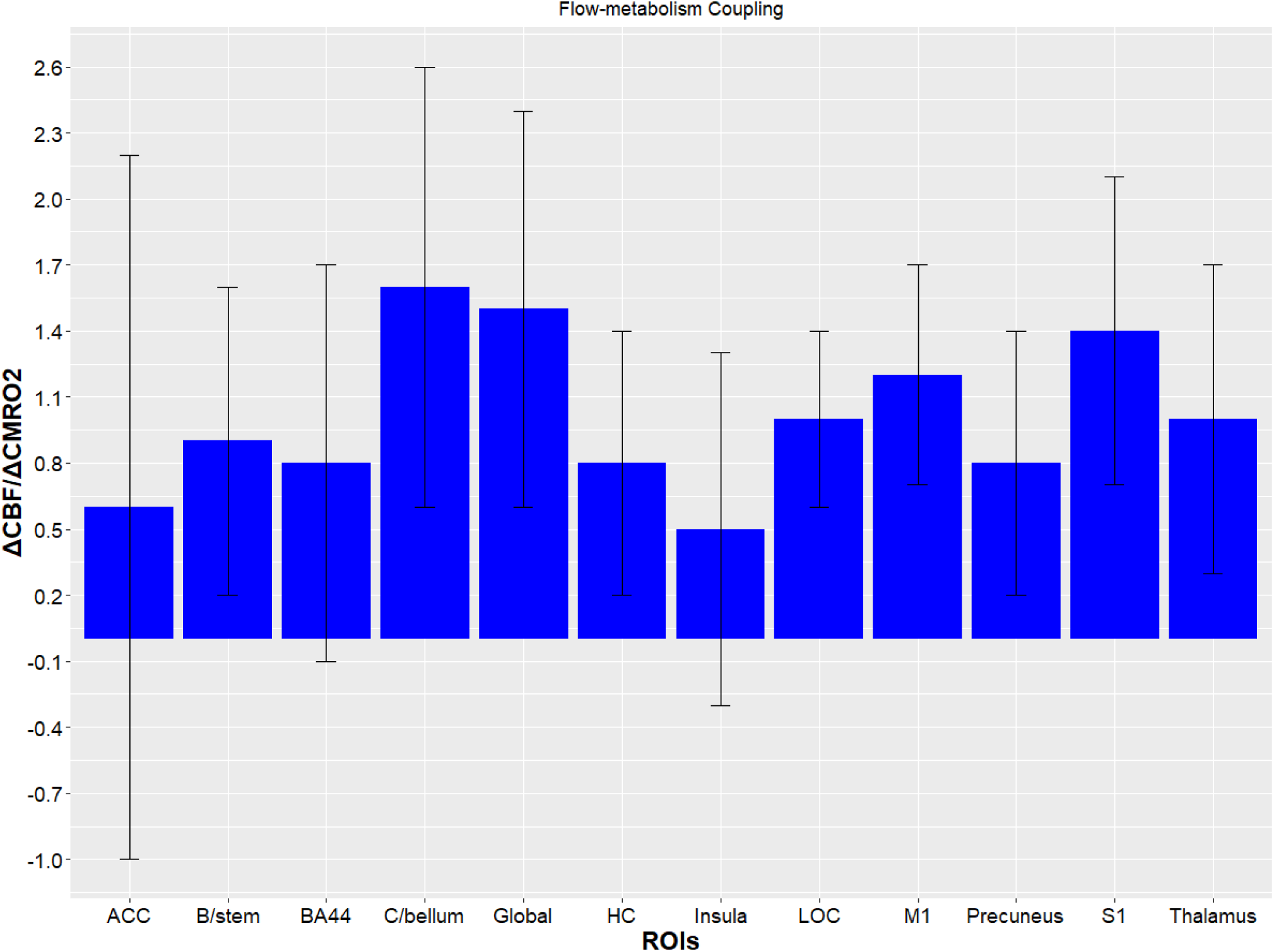
Mean coupling ratios (ΔCBF/ΔCMRO_2_) all SRT task blocks calculated as a per subject average, SEM represents SEM across subjects. B/stem = brainstem, C/bellum = Cerebellum, HC = Hippocampus.

Paired t-tests were carried out to identify significant changes between the first and second, and first and last blocks. Significant CMRO_2_ changes between blocks were observed in the global task ROI, the cerebellum and between the first and last blocks in M1 and S1. Significant BOLD changes between blocks were observed in most regions, except BA44, and the thalamus between block 1 and block 6. This was also the case for CBF where changes were not observed in BA44, the brainstem, hippocampus and precuneus between block 1 and 2, and the thalamus between block 1 and 6 (see figure 5A-K for a summary of statistically significant changes for each ROI and block.

Outliers in the BOLD and CBF and CMRO_2_ data were identified using Tukey’s method(Tukey, 1977) where the interquartile range is multiplied by 1.5 to define a reasonable range. A number of subjects were then rejected from each ROI on this basis (see supplementary table 1).

### Flow Metabolism Coupling

The flow-metabolism coupling ratio, *n*, was investigated in each region comparing differences between sequence block 1 and block 2 and between sequence block 1 and block 6 using repeated measures ANOVAs. Figure 6 shows the mean coupling across each ROI. Despite observed differences, there were no statistically significant differences in coupling between any ROIs or timepoints. Of the 12 ROIs, global, M1, S1, cerebellum and the hippocampus showed what could be considered ‘typical’ coupling. The precuneus and brainstem displayed typical coupling until the final 2 blocks (Figure 5A-F). The LOC, insula, BA44, ACC, and thalamus had more atypical coupling relationships (see figure 5G-K) where the BOLD response was positive despite a negative CMRO_2_ change from baseline. Typical coupling, in this instance refers to ROIs where all BOLD, CBF and CMRO_2_ follow the same pattern, e.g. sustained responses above baseline in all parameters. An example of atypical coupling is in the insula where there was a sustained BOLD task response across the task yet CBF and/or CMRO_2_ was near or below baseline levels.

### FMRI and Behaviour

Regression analysis was conducted to determine whether RT was predicted by BOLD, CBF or CMRO_2_. For this, we focused on the first and last SRT task blocks. Multiple linear regression was carried out across subjects for each ROI with RT as the dependent variable, and BOLD, CBF and CMRO_2_ as independent variables. Neither BOLD, CBF nor rCMRO_2_ were predictors of RT at the beginning or end of the task with all regression coefficients below 0.2, and p-values < 0.05 (data not shown).

## Discussion

In this study we have shown changes from baseline in BOLD, CBF and CMRO_2_ in multiple brain regions recruited during performance of an SRT task, as well as changes in the amplitude of these responses from early to late task blocks. The pattern of changes was heterogeneous across regions, but the largest response amplitudes occurred in the first task block. Outside of M1 and S1, in areas where a statistically significant CMRO_2_ response was detected in early blocks (BA44, precuneus and cerebellum), CMRO_2_ returned to within baseline levels by the third task block. In contrast, BOLD responses were sustained (i.e. significantly different from baseline) across blocks in most ROIs except for the precuneus and hippocampus in later blocks. In the cerebellum, a sustained CBF response was also observed in the absence of a significant CMRO_2_ increase after the first sequence block.

In random task blocks, only M1 and S1 showed significant CBF and CMRO_2_ responses, however CBF responses across all 3 random blocks were observed in BA44 and the cerebellum, and in random block 1 in the ACC, precuneus and LOC.

The mean flow-metabolism coupling ratio *n* over all sequence blocks in the cerebellum was 1.6, which was over 3 times that of the insula where the coupling ratio was 0.5 (see figure 6). Despite this, coupling was not statistically different between any ROIs, or between task blocks within ROIs, perhaps as inter-subject variability was high.

The increase in RT for random task blocks was not reflected by increases in the BOLD or CBF responses, and RT in SRT blocks did not correlate significantly with the imaging data, despite fMRI responses and RT both decreasing in a similar fashion across blocks. Therefore, the BOLD, CBF or CMRO_2_ responses cannot be assumed to reflect sequence specific learning directly. What we are demonstrating, is the energetic adaptations occurring with repeated execution of a motor sequence, and therefore this study has important implications for the study of plasticity and higher cognitive processes such as learning. With appropriate tasks in future studies, CMRO_2_ could be calculated during early and late phases of learning as well as before and after behavioural or drug interventions to study energetic changes during learning and memory processes, and to evaluate the neural changes brought out by interventions, as CMRO_2_ is assumed to be closely coupled to neuronal activity.

### Energetic Adaptations during the SRT

Task responses and response reductions over time were identified in regions typical of a visuomotor learning task; M1, S1, the ACC, cerebellum, insula and LOC, areas involved in motor performance, visual processing, attention and information processing. The cerebellum is critically involved in motor learning and coordination of voluntary movement, therefore decreases in this area may represent a shift towards automated performance of the motor task, given the sustained CBF response and CMRO_2_ near baseline levels. The insula has been associated with hand to eye motor coordination, while the LOC is primarily involved in shape discrimination; signal reductions in these regions, combined with the absence of a significant CMRO_2_ increase during task performance may also be attributed to rapid task adaptation and low difficulty level of the task.

Outside of M1 and S1, key regions involved in motor execution, several ROIs including the ACC, thalamus and BA44 showed a sustained BOLD response in the absence of a significant CMRO_2_ response. BOLD signal increases without notable CMRO_2_ responses may be due to signalling mechanisms leading to a preparatory haemodynamic response (Buxton, 2010; Buxton, Griffeth, Simon, & Moradi, 2014; Gjedde, Vafaee, & Gjedde, 2004) which, due to the low task difficulty, is not subsequently required as only a small CMRO_2_ increase is needed to perform the task. Other regions such as the insula, LOC and hippocampus showed CMRO_2_ reductions from baseline in later blocks, with only the hippocampus also showing BOLD reductions from baseline, however these CMRO_2_ changes were not statistically significant. Low CMRO_2_ responses in regions where the BOLD, and sometimes CBF, response was sustained across blocks, such as in cerebellum, ACC, thalamus and BA44, demonstrates the importance of acquiring metabolic measures in addition to the BOLD response. Taking BA44 as an example, the BOLD response alone suggests a significant BOLD increase was sustained for all sequence and random blocks. However, the CBF data shows only significant changes for the first sequence blocks and all random blocks, and a CMRO_2_ increase was detected only in the first task block. BA44 has been previously associated with motor adaptation (Shannon et al., 2016), where reductions in oxygen consumption and functional connectivity with V1 were observed following a motor adaptation task. The CBF and CMRO_2_ data suggest that motor adaptation occurred quickly, with greater effort required for the first sequence block and random blocks. The sustained BOLD may be a result of the preparatory motor response (Gjedde et al., 2004) with only a small, or negligible, subsequent metabolic increase.

As we did not measure aerobic glycolysis, it is not possible to identify changes in both this and CMRO_2_. However, increases in aerobic glycolysis may also explain the absence of CMRO_2_ increases during the task in regions such as BA44, if there was increased aerobic glycolysis and reduced oxygen metabolism with task adaptation (Shannon et al., 2016). Rather than a small increase in metabolic demand during the task, it could also be that there is a shift towards greater aerobic glycolysis, hence the lack of an observed CMRO_2_ increase in many ROIs, including BA44, after the initial sequence block. The sustained CMRO_2_ response observed in M1 and S1 may be explained by ongoing recruitment of these regions for motor execution, rather than adaptation or learning, processes may rely more on increases in aerobic glycolysis. As shown in figure 5A-K, the greatest responses in BOLD, CBF and CMRO_2_ are always observed in the first SRT block. This is likely to be a result of the initial demands of the task, focusing visual attention on the task and selecting the correct motor responses, as well as processing task information to identify the repeating sequence. In this particular task, subjects quickly adapted to the task demands, as evidenced by the rapid improvement in accuracy and reduction in response time. This fast adaptation may explain the sharp reduction in BOLD and CBF responses, as well as the small, on average, CMRO_2_ increases from baseline in later blocks, as this adaptation led to a reduced neuronal demand.

While the neurobiological phenomena discussed above may explain the data, it must also be kept in mind that the CMRO_2_ measurements obtained using calibrated fMRI are prone to noise and therefore lower power than individual optimised measurements of each parameter. CMRO_2_ measured using calibrated fMRI being derived from BOLD and ASL CBF signals suffers from a lower contrast-to-noise ratio than BOLD, limiting the detectability of small CMRO_2_ changes. As a result, changes from baseline may not have been detected, whereas BOLD and CBF changes, more robust over short timescales, were detected.

Signal reductions over task blocks observed in this study have been reported previously for BOLD and PET (Floyer-Lea & Matthews, 2004; Laguna & Cognitive, 1995) with the largest reduction also occurring between the first and second blocks. Fernandez-Seara et al. (2016) used ASL to investigate CBF changes during two separate 6-minute explicit learning tasks where a different pattern of sequential finger movements was trained in each task. To investigate learning-related changes, 3 learning phases were defined representing early to intermediate learning. CBF decreases relative to CBF measured during a control block were reported bilaterally for regions recruited during task performance, and perfusion reached levels comparable to baseline by the final task block in agreement with many ROIs in the current study. However, in contrast to the current study, where no perfusion increases over time were found, Fernandez-Seara et al. (2016) found perfusion increases with task practice in somatosensory cortex, the posterior insula and putamen, cingulate cortex and left hippocampus which may be due to differences in the task design or ASL sequence, as PCASL, which provides increased perfusion signal was used by Fernandez-Seara et al. (2016).

Although significant changes in CBF and CMRO_2_ without accompanying BOLD changes were not typically observed, acquiring CBF and CMRO_2_ data is still valuable as it can provide an insight into the main processes contributing to the measured BOLD signal. This is evident where BOLD and CBF remain elevated from baseline across the task despite CMRO_2_ being close to, or reduced from, baseline. Most regions showed similar patterns at the beginning of the task, significant BOLD and CBF responses, accompanied by CMRO_2_ changes in motor regions, and the cerebellum, and a sharp drop in signal by block 2. In agreement with Buxton et al. (2014), the CMRO_2_ response suggests more rapid adaptation to the task than the BOLD or CBF data; it is possible that neural adaptation occurs faster than the haemodynamic response adaptation. Changes in later blocks became more variable across ROIs, but overall CMRO_2_ responses were close to baseline, in contrast to the larger responses seen in the first SRT block in many ROIs. This pattern may be similar to previously reported rapid adaptation followed by slower, more gradual adaptations which are likely to continue beyond the short task duration (Doyon & Benali, 2005).

### Flow-metabolism Coupling

In this study, no significant differences in flow-metabolism coupling were observed between ROIs or task blocks, despite group mean coupling ratios ranging from 0.5-1.6 across ROIs, therefore speculation about the differences observed must be limited. Observed values were lower than coupling ratios previously reported in the visual cortex during visual stimulation of 2.2 (Uludağ et al., 2004) and ~1.8 (Kim et al., 1999), this may derive from weaker visual stimulation in the present study compared to checkerboard stimuli commonly used to activate V1. Ratios of change in CBF to change in CMRO_2_ of 3:1 have been reported for graded motor activation from 0.5-3hz (Kastrup, Kru, Neumann-Haefelin, Glover, & Moseley, 2002) and in M1, a group average value of ~3.4 has been reported during finger-thumb opposition (Chiarelli et al., 2007). This is much larger than the changes observed in the current study, but the values reported for the current work are comparable to values of 1.6 reported for the medial temporal lobe during a memory encoding task (Restom, Perthen, & Liu, 2008) which required motor responses to images presented in blocks of familiar and unfamiliar stimuli. The variation across studies is likely to be due to differences in the type of stimulation or task used, but also reflect differences in neurovascular coupling across the brain. The lack of statistically significant differences between blocks or regions in the current study may be explained by the large inter-subject variation, both between regions and blocks (see figure 6).

Typical coupling between CBF and CMRO_2_ in brain regions outside of visual, motor and somatosensory cortices has not been clearly established, nor is it clear how processes such as skill acquisition affect the coupling ratio *n*. Uludağ et al. (2004) reported that coupling ratios during activation and deactivation (due to task-related signal decreases) are similar in the visual cortex, but it is not clear if this trend is uniform across the brain and under different levels of cognitive load. As mentioned in the introduction, changes in coupling may occur as a result of changes in aerobic glycolysis as less glucose is metabolised through oxidative phosphorylation during and following learning (Shannon et al., 2016). In visuomotor adaptation and learning tasks, the coupling ratio *n*, and changes in *n* with task practice, may represent the LTP and LTD processes involved in learning, and which underlie plasticity in the healthy brain. However, additional studies which track CBF and CMRO_2_ after task performance are needed to establish whether energetic changes suggestive of LTP and LTD can be identified using fMRI during short motor adaptation tasks.

Finally, brain measures did not correlate significantly with gains in performance in the form of accuracy and reaction time. Contributing to this observation may the low level of difficulty such that ceiling effects were observed in the performance data, alongside very low variability in performance across subjects. Coupled with the high variability of the brain data, this may explain why no correlations were observed. Therefore, although motor adaptations were observed in the form of performance improvements and signal reductions, the signal changes cannot be directly interpreted as sequence specific motor learning. In the pseudorandom task blocks performance was lower, however, there was no significant difference in BOLD or CBF in these blocks compared to the SRT tasks before and after (data not shown). This suggests that the changes in BOLD, CBF and CMRO_2_ were not due to sequence specific motor learning but more to adaptation to repeated finger movements.

### Limitations

The principal limitation is the low contrast-to-noise of CMRO_2_ estimates, largely arising from low CNR of ASL CBF measurement. Previous work studying cerebrovascular changes during cognitive tasks has used much longer paradigms than the 12-minute task used here, with some exceptions (Fernandez-Seara et al., 2016b). Only 6 sequence blocks were included here due to time limitations within the scan session, meaning that SNR was relatively low for the purposes of looking at individual 45 second block activity. Additional blocks would allow ‘chunking’ of the data into early, intermediate and late blocks to examine differences between the 3 stages of learning with increased SNR. For example, Olson et al. (2006) incorporated three 20-minute SRT blocks in their protocol which served to dilute the effects of transient anomalies in activity and build a more reliable picture of average group responses. While the 12-minute task is useful for measuring short term adaptation differences between patients and controls and to limit head motion and fatigue effects, to quantitatively assess motor learning, longer paradigms should be used along with larger sample sizes to establish whether the variability observed in this study is due to real individual differences or simply the low CNR inherent in ASL data.

### Conclusions

This study demonstrates the use of calibrated fMRI to detect regional BOLD, CBF and CMRO_2_ responses to a short motor adaptation task, and the changes in the amplitude of these responses over time in a sample of healthy adults. The most interesting result of this study is the finding that in several regions including the ACC, BA44, thalamus and cerebellum, there was a sustained BOLD haemodynamic response in the absence of a significant CMRO_2_ increase from baseline which suggests that the brain may continue to elevate the supply energy even after actual utilisation (CMRO_2_) has reduced to near baseline levels. Therefore, relying on BOLD data alone in behavioural studies can mask the nature of underlying metabolic responses and their changes over time with repeated task performance. With refinements to the task and MR acquisition, calibrated fMRI could be used to study energetic changes during learning in the healthy brain and to investigate the vascular and metabolic changes underlying reduced cognitive and motor function and limited plasticity in ageing and disease.

## Supplementary Data

**Table S1.** Number of outliers removed from analysis of each ROI

| ROI | Number of outliers removed from imaging data |
| --- | --- |
| Global | 1 |
| ACC | 3 |
| BA44 | 2 |
| Brainstem | 0 |
| Cerebellum | 2 |
| Hippocampus | 1 |
| Insula | 1 |
| LOC | 1 |
| M1 | 1 |
| S1 | 3 |
| Precuneus | 3 |
| Thalamus | 3 |

## Acknowledgements

The authors would like to thank Dr. Claudine Gauthier and Dr. Christopher Steele for helpful discussions on data analysis and interpretation and Dr. Ilona Lipp for assistance with analysis pipelines and task development. RGW and VT are supported by the Higher Education Funding Council for Wales.

